# Object category warps the topography of object space for behavior

**DOI:** 10.64898/2026.03.11.711098

**Authors:** Xinchi Yu

## Abstract

Recent neuroimaging work suggests that the primate ventral visual stream is organized according to a 2D object space, defined by two axes, namely animate-inanimate and stubby-spiky. While this 2D geometry explains substantial neural variance, does it serve as a static, category-independent scaffold for behavior? We addressed this question using a large-scale visual foraging dataset in humans (*n* = 511). By employing a rank order analysis and a novel reverse-correlation-inspired “landscape analysis” to map behavioral preferences in visual foraging “back to” the object space, we revealed a critical dissociation. We found that the behavioral preference topography in this space is highly stable within object categories, yet it fails to generalize across categories, frequently exhibiting reversed polarities. These findings challenge a “universal topography” hypothesis where the 2D object space guides behavior regardless of object category. Instead, our results demonstrate that object categories warp the topography of the 2D object space, supporting a “category-dependent topography” view. Taken together, our current results raise the question of how the object space is flexibly remapped in support of human behavior.

---

A pioneering macaque neuroimaging study by Bao and colleagues revealed that the inferotemporal cortex is organized into four sets of patches defined by two axes: animate-inanimate and stubby-spiky ^1^. This 2D “object space” can be effectively modeled by different subspaces in the first two principal components (PC1-2) of deep neural network (DNN) representations, derived from applying PCA to a larger set of visual objects ^1–4^. This object space, signatured by the two representational axes, has been broadly replicated in human neuroimaging ^2,4^. A critical question remains: Does the topography of object space serve as a static, universal scaffold for human visual behavior?

To answer this question, we leveraged a large-scale human visual foraging dataset (*n* = 511) ^3^. Ten real-world object categories were involved in the study (e.g., birds, hands, mattresses), spanning four “quadrants” across animate-inanimate and stubby-spiky: (animate, stubby), (animate, spiky), (inanimate, stubby), (inanimate, spiky). Note that here “(in)animate” serves as a shorthand for (in)animate-looking, as objects can visually fall into the animate-looking quadrants independent of their conceptual animacy. To match the number of object categories for each quadrant, here we only analyze data for eight out of ten categories, with two categories per quadrant (**Figure 1B**; another selection of eight categories rendered qualitatively same results, see **Supplementary Materials**). In each trial, participants search for a target object (**Figure 1A**). Multiple instances of the same target object are presented on the screen along with distractors from the same object category. Behavioral performance for each trial was characterized as the number of correct clicks per second, where more correct clicks second^−1^ signifies better performance.

**Figure 1.**
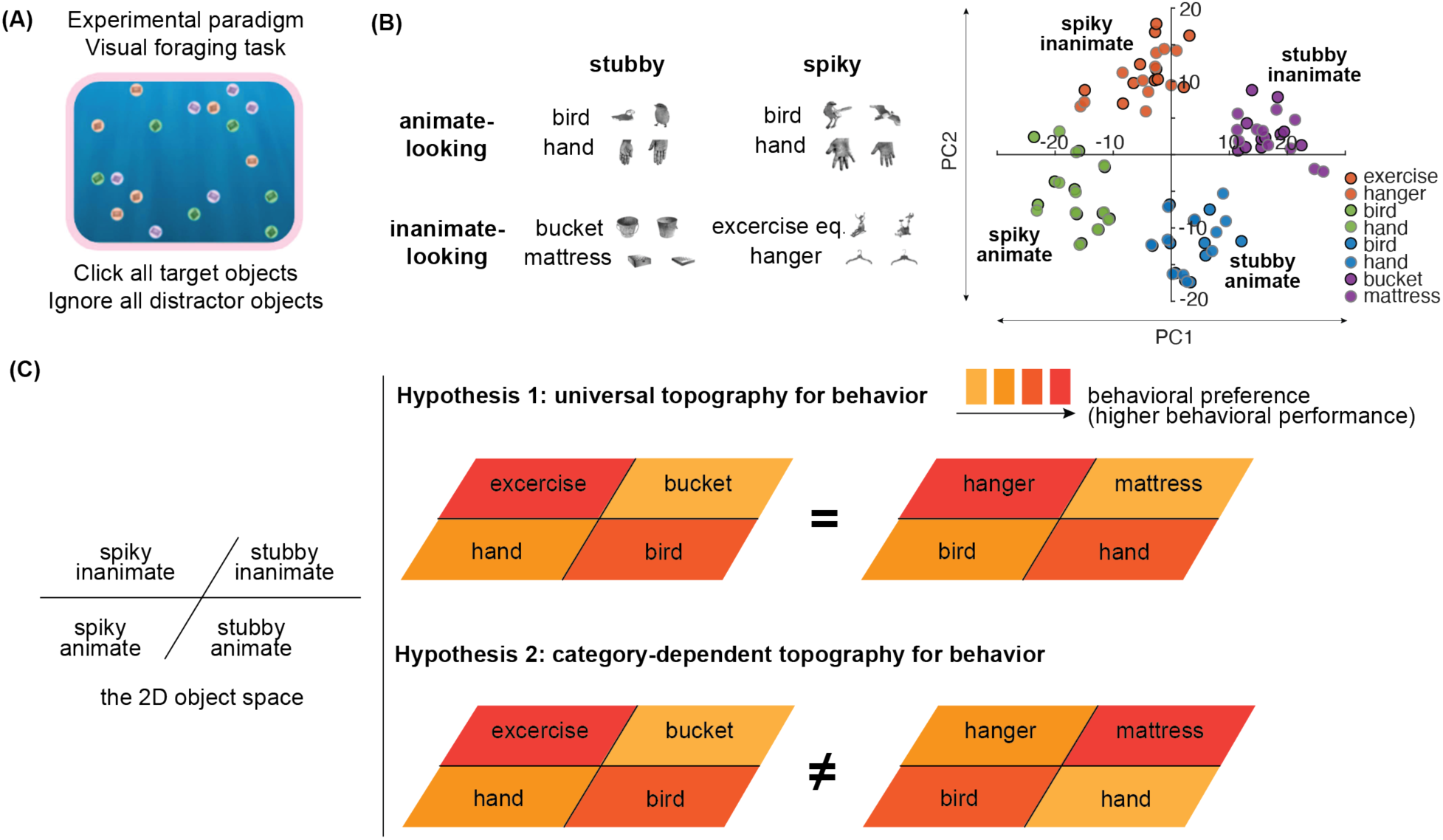
**(A)** Illustration of the experimental paradigm: the visual foraging task. **(B)** (Left Panel) Eight categories spanning the four quadrants that entered our analysis, with two example exemplars each. (Right Panel) All exemplars’ locations in the 2D object space. exercise eq. or exercise stands for “exercise equipment”. **(C)** Two competing hypotheses on how the object space maps onto behavior. The “universal topography for behavior” posits that the topography of the object space dictates behavioral performance regardless of object categories. Conversely, the “category-dependent topography for behavior” hypothesis posits that the topography of the object space is category-dependent.

We contrasted two competing hypotheses regarding how the object space maps onto behavior (**Figure 1C**). The “universal topography for behavior” hypothesis posits that the topography of the object space dictates behavioral performance regardless of object categories. This is consistent with a view where pre-determined object category does not play a unique role in neural representation and behavior ^5^. Conversely, the “category-dependent topography for behavior” hypothesis posits that the topography of the object space is category-dependent. This is consistent with a view suggesting that feature and category play non-identical roles in neural representation and behavior ^6^.

## RESULTS

### Rank order analysis reveals within-category consistency of object space topography

First, we want to test whether people have consistent behavioral preferences across trials, when we select four object categories comprising one category from each of the four quadrants. As every trial involved non-overlapping targets, this tests the extent to which the topography of behavioral preference generalizes across different exemplars of the same object categories. This is therefore an analysis on within-category consistency. To this end, we conducted a rank order analysis, where for each iteration we sample trials randomly into two non-overlapping split-halves. We then computed the mean behavioral performance for each of the four object categories within each half. After this, we quantified the consistency between the rank order of behavioral preferences observed in the first half and that in the second half, with Spearman’s ρ (**Figure 2A**). As a baseline, we also computed the Spearman’s ρ between the rank order of the first half and a shuffled order of the second half (ρ_shuffled_). By repeating this procedure across 1,000 iterations, we tested whether the rank order of behavioral preferences established in one set of exemplars was reliably preserved in an independent set.

**Figure 2.**
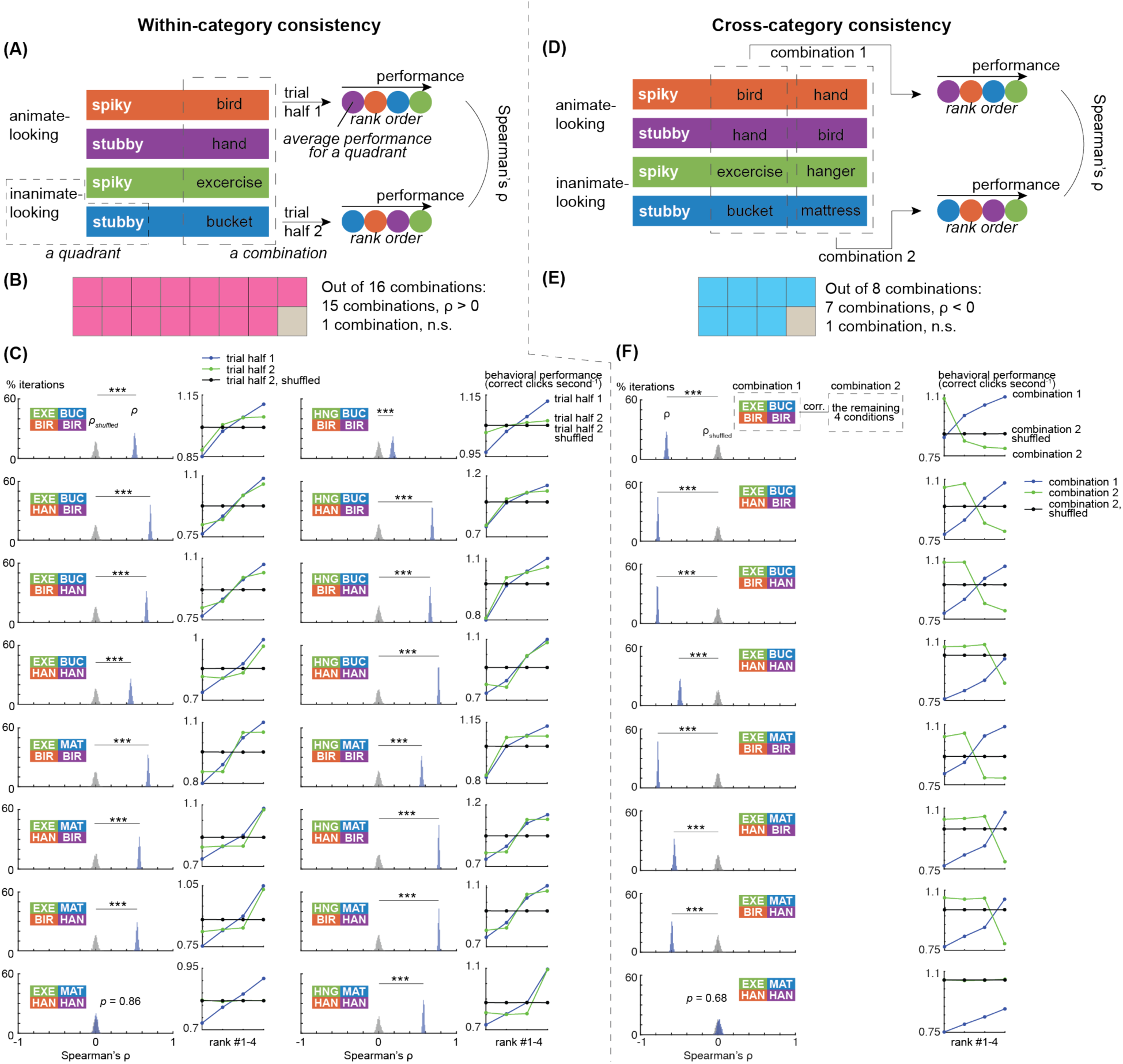
Results for the rank order analysis, where (A)-(C) are on within-category consistency, and (D)-(F) are on cross-category consistency. Abbreviations: EXE: exercise equipment; BUC: bucket; BIR: bird; HAN: hand; MAT: mattress; HNG: hanger. **(A)** Illustration of the procedure of within-category consistency analysis. **(B)** Summary of the results for the within-category consistency analysis. Pink blocks stand for statistically significant positive results, and gray blocks stand for statistically non-significant results. n.s.: non-significant. **(C)** Left panels for each pair of plots: Bootstrapped distribution of Spearman’s ρ (blue) and the shuffled null distribution (gray), for a certain selection of four categories (in abbreviation). Right panels for each pair of plots: The blue dots reflect behavioral performance, ranked for a split half of trials (trial half 1). The green ones show behavioral performance of the remaining half (trial half 2), ranked by data from the first half (both averaged across 1,000 iterations). The black dots show an estimation of chance, by shuffling the rank orders of the second half. This visualization procedure is similar to other studies on split-half consistency ^13,14^. ***: *p* < 0.001. **(D)** Illustration of the procedure of cross-category consistency analysis. **(E)** Summary of the results for the within-category consistency analysis. Blue blocks stand for statistically significant negative results, and gray blocks stand for statistically non-significant results. n.s.: non-significant. **(F)** Left panels for each pair of plots: Bootstrapped distribution of Spearman’s ρ (blue) and the shuffled null distribution (gray), for a certain selection of four categories (in abbreviation). Right panels for each pair of plots: The blue dots reflect behavioral performance, ranked for a split half of conditions (combination 1). The green ones show behavioral performance of the remaining half (combination 2), ranked by data from the first half (both averaged across 1,000 iterations). The black dots show an estimation of chance, by shuffling the rank orders of the second half. ***: *p* < 0.001.

We found that the rank order of behavioral preference across the four quadrants is reliably consistent, for a specific combination of categories across the four quadrants. For 16 combinations of four categories, 15 out of 16 combinations had a Spearman’s ρ significantly larger than the shuffled distribution, exact two-tailed *p’s* < 0.001 (**Figure 2B,C**). Spearman’s ρ was not significantly different from the shuffled distribution for the remaining one combination (*p* = 0.86).

### Rank order analysis reveals a lack of cross-category consistency of object space topography

Then we want to test whether people have consistent preferences across categories, when we compare across four object categories comprising one category from each of the four quadrants against the remaining four object categories. To this end, we partitioned the eight object categories into two non-overlapping sets, each comprising one distinct category from each of the four quadrants. We then computed the rank order of behavioral performance across the four quadrants for the first set and correlated it with that of the second set, with Spearman’s ρ (**Figure 2D**). The above procedure was repeated across 1,000 iterations by resampling the participants with replacement. This applied for all 8 pairs of possible combinations of the object categories.

In stark contrast to the within-category analysis, we found no evidence for a universal topography for behavioral preference across categories. Rather, most conditions showed a reliable inverse correlation across the two combinations of categories. For 8 combinations of four categories, 7 out of 8 combinations had a Spearman’s ρ significantly lower than a shuffled distribution, exact two-tailed *p’s* < 0.001 (**Figure 2E,F**). Spearman’s ρ was not significantly from the shuffled distribution for the remaining one combination (*p* = 0.68).

### Developing and validating the landscape analysis

One remaining question is the extent to which the Euclidean distance between the target and distractor in the object space differ on a trial-by-trial basis. As target-distractor distance does influence visual foraging performance ^7^, we need a method that can take this into account. To address this concern, we devise a novel “landscape analysis”, inspired by the reverse correlation technique ^8,9^, that enables us to parse out effects of target-distractor distance at a trial-by-trial basis.

The core idea is to recover the behavioral performance across all pixels in the object space based on a certain number of trials, by computing, for each pixel, the partial correlation between behavioral performance and the target’s proximity to that pixel, while statistically controlling for the trial-specific target-distractor distance (**Figure 3A**; see **Methods**). The governing logic is that, if a pixel is more behaviorally preferred, the closer the target is to this pixel, the better the behavioral performance. Following this idea, a pixel’s behavioral performance is characterized as the opposite value of the abovementioned partial correlation coefficient, as behavioral performance should increase with the decrease of pixel-target distance. Through ground-truth simulations (**Figure 3B-D**), we confirmed that this method succeeds in recovering the spatial direction in which behavioral performance increases/decreases (for details of the simulation, see **Methods**).

**Figure 3.**
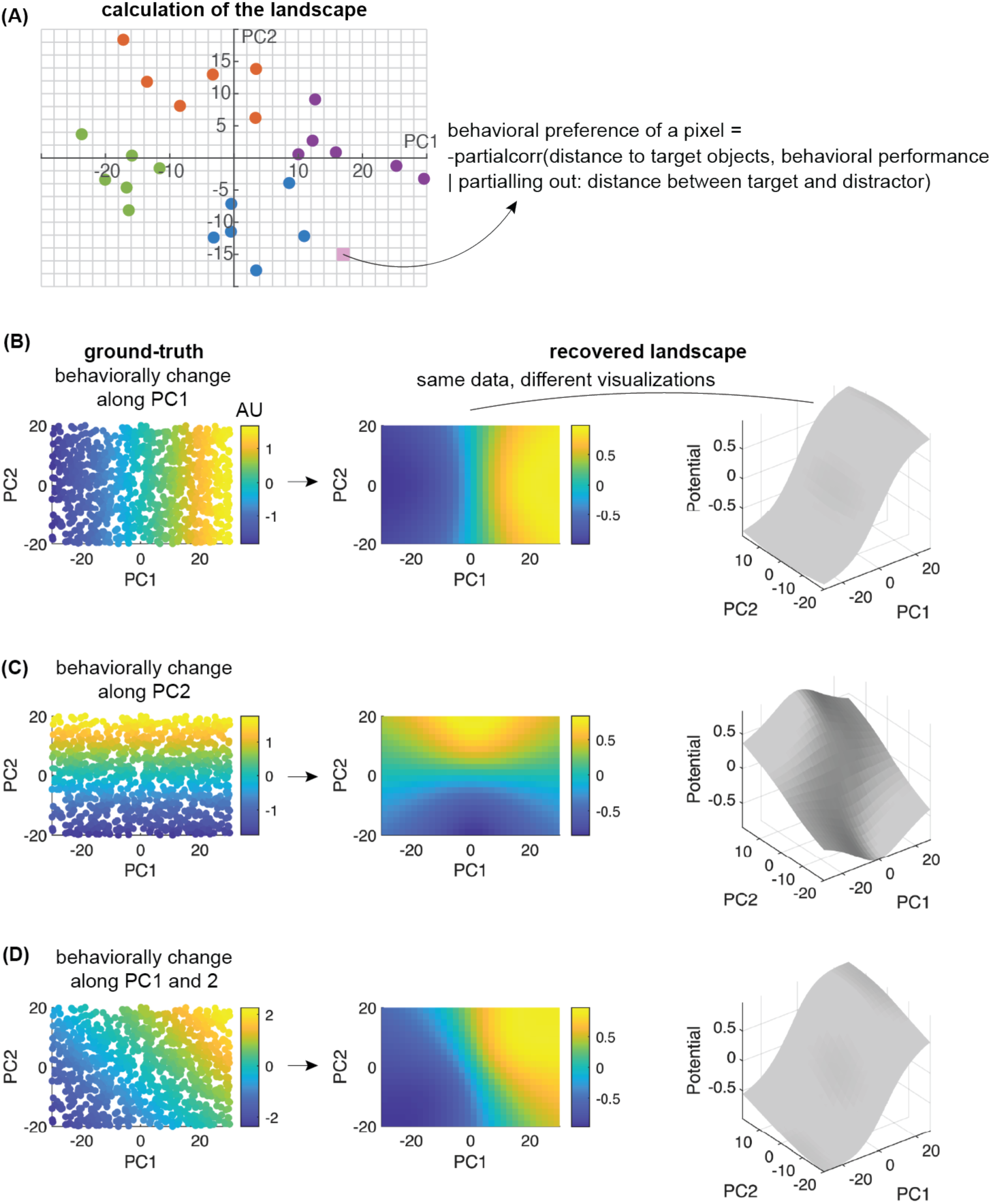
Illustration of the procedure of landscape analysis (A), and validation of this analysis (B-D). **(A)** Illustration of how behavioral preference for each pixel is calculated, with a certain selection of trials (circles). **(B)** The leftmost panel shows sampled 1,000 trials from a ground-truth where behavioral performance (in arbitrary units, AU) increases along PC1. Behavioral performance for each trial is color-coded. The recovered landscape is calculated following the procedure in (A), and is visualized with colors or heights in the middle and the rightmost panels respectively. **(C)** Similar to (B), but the ground-truth is that behavioral performance increases along PC2. **(D)** Similar to (B), but the ground-truth is that behavioral performance increases along the diagonal.

### Landscape analysis corroborates within-category consistency of object space topography

Similar to the rank order analysis, we want to test whether people have consistent behavioral preference landscapes across trials, when we select four object categories comprising one category from each of the four quadrants. The procedure was in line with the rank order analysis, replacing rank order across iterations to the rank order of all pixels in the landscape. For each iteration, we sample trials randomly into two non-overlapping split-halves. We then computed the behavioral preference landscape for each of the four object categories within each half. After this, we quantified the consistency between the landscape observed in the first half and that in the second half, with Spearman’s ρ (**Figure 2A**). As a baseline, we also computed the Spearman’s ρ between the rank order of the first half and a shuffled order of the second half. By repeating this procedure across 200 iterations (fewer iterations due to increased computational cost), we tested whether the landscape established in one set of exemplars was reliably preserved in an independent set.

Controlling for trial-by-trial target-distractor similarity with the landscape analysis, we again found that the behavioral preference landscape across the four quadrants is reliably consistent, for a specific combination of categories across the four quadrants. Trial-by-trial target-distractor similarity was quantified as the Euclidean distance between the target and the distractor in the 2D object space, as derived in the original study ^3^. For 16 combinations of four categories, 15 out of 16 combinations had a Spearman’s ρ significantly larger than a shuffled distribution, exact two-tailed *p’s* < 0.001 (**Figure 4B,C**). Spearman’s ρ was significantly lower than the shuffled distribution for the remaining one combination, but with a small numerical value that is hardly meaningful (*ρ_mean_* = -0.06, *p* < 0.001).

**Figure 4.**
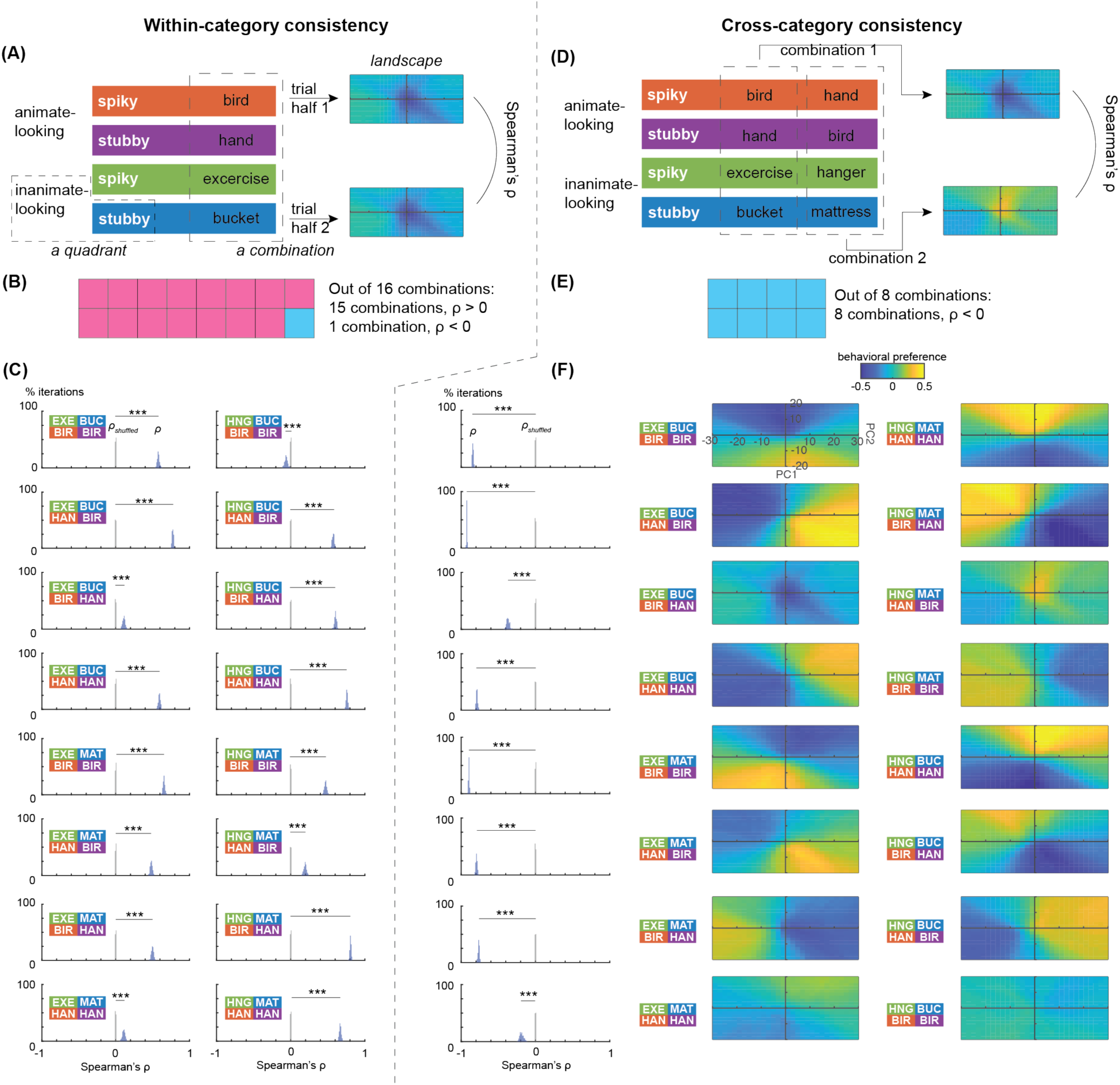
Results for the landscape analysis, where (A)-(C) are on within-category consistency, and (D)-(F) are on cross-category consistency. Abbreviations: EXE: exercise equipment; BUC: bucket; BIR: bird; HAN: hand; MAT: mattress; HNG: hanger. **(A)** Illustration of the procedure of within-category consistency analysis. **(B)** Summary of the results for the within-category consistency analysis. Pink blocks stand for statistically significant positive results, and blue blocks stand for statistically significant negative results. **(C)** Bootstrapped distributions of Spearman’s ρ (blue) and the shuffled null distribution (gray), for a certain selection of four categories (in abbreviation). ***: *p* < 0.001. **(D)** Illustration of the procedure of cross-category consistency analysis. **(E)** Summary of the results for the within-category consistency analysis. Blue blocks stand for statistically significant negative results. **(F)** The left most column of panels: Bootstrapped distribution of Spearman’s ρ (blue) and the shuffled null distribution (gray), for a certain selection of four categories (in abbreviation). On the right of each panel are the averaged landscapes for the two corresponding combinations. ***: *p* < 0.001.

### Landscape analysis corroborates a lack of cross-category consistency of object space topography

Similar to the rank order analysis, we want to test whether people have consistent preferences across categories, when we compare across four object categories comprising one category from each of the four quadrants against the remaining four object categories. The procedure was also in line with the rank order analysis. We partitioned the eight object categories into two non-overlapping sets, each comprising one distinct category from each of the four quadrants. We then computed the behavioral preference landscape across the four quadrants for the first set and correlated it with that of the second set, with Spearman’s ρ (**Figure 4D**). The above procedure was repeated across 200 iterations by resampling the participants with replacement. This applied for all 8 pairs of possible combinations of the object categories.

Again, in stark contrast to the within-category analysis, we found no evidence for a universal topography for behavioral preference across categories. Rather, all 8 conditions showed a reliable inverse correlation across the two combinations of categories, exact two-tailed *p’s* < 0.001 (**Figure 4E,F**).

## DISCUSSION

Leveraging a large-scale visual foraging dataset (*n* = 511) and employing complementary rank order analysis and a novel “landscape analysis”, we uncovered two properties of the object space. First, we found that the landscape (i.e., topography, geometry) for behavioral preferences is highly stable within object categories. Second, and crucially, we observed that this landscape does not generalize across different object categories.

First, our results extend prior neurophysiological observations by demonstrating that the 2D object space is not merely a property for neural representations ^1,2,4^, but is also a robust model for human behavior. Second, our results reveal critical limitations of a “universal topography for behavior” view of the object space, where the object space serves as a static and category-independent scaffold for behavior (**Figure 1C**). Our results instead suggest that the topography of the object space can depend on the categories selected. This finding extends and complements the observations in the original study ^3^. In their pioneering work, Sigurdardottir and Ólafsdóttir uncovered positive associations between task performance for faces in a certain quadrant and non-face objects in the same quadrant, suggesting a certain extent of generalization of object space topography across object categories. Our results with eight non-face object categories revealed the boundary conditions of this generalization: we found that the correspondence between the 2D object space and behavior does not always generalize well across object categories. Uncovering the determinants of the extent of generalization is therefore a curious direction of future research. Third, our newly-devised “landscape” analysis offers a new lens to study the object space, and allows us to easily parse out trial-by-trial confounds. Typical reverse-correlation studies correlate behavioral outcomes back to the stimuli to uncover the pixels critical for a certain task outcome ^9^. Our landscape analysis instead correlates behavioral outcomes back to an abstract object space. As reverse-correlation enables *generating* stimuli that maximizes a certain behavioral outcome ^10^, our landscape analysis paves the way to generating objects with better behavioral performance.

Taken together, our results support a view where behavior is explained by a combination of feature space and object category ^6,11,12^. Moving beyond mono-cause models for behavior, our current study highlights how human behavior emerges from dynamic interactions between a universal feature space and higher-level object categories.

## Acknowledgements

We would like to thank Heida Maria Sigurdardottir and Inga María Ólafsdóttir for extensive comments and suggestions.

## Author contributions

X. Y. conceptualized the analyses, performed the analyses, and wrote the manuscript.

## Data and code availability

Analysis scripts will be openly shared on OSF upon acceptance. Raw data are openly shared by the authors of ref. ^3^ (https://osf.io/2jn6c).

## EXPERIMENTAL AND PARTICIPANT DETAILS

### Participants

The study utilized a large-scale, openly-available visual foraging dataset ^3^. For a comprehensive report of participants and the procedures please refer to the original article. This dataset consists of data from 511 participants. 357 were women, 144 were men, and 10 were nonbinary, other gender, or chose not to answer. 25 people were between the ages of 18-24, 86 were 25-34 years, 134 were 35-44 years, 132 were 45-54 years, and 134 were 55 years of age or older. All participants provided informed consent in accordance with the ethical guidelines of the original study.

### Defining the object space

The object space was defined with a reference set of 120 animals and 120 inanimate objects spanning a variety of categories. The object space was defined as the first two principal components (PC1 and PC2) of the second fully connected (fc) layer of the VGG-16bn deep neural network (DNN). We used the object space coordinates for each stimuli object provided by ref. ^3^, which were derived from projecting the DNN representation onto the object space.

### Visual foraging stimuli and task

As our current study focuses on object categories, and that the original study involved multiple face conditions within the same quadrant, for the current study we only consider conditions that involved real-world, non-face objects. This leaves a total of 10 categories of real-world objects (e.g., birds, hands, buckets, mattresses). There were two categories for the (spiky, inanimate) quadrant: hangers, exercise equipment. There were two categories for the (stubby, inanimate) quadrant: mattress, buckets. There were two categories for the (spiky, animate) quadrant: birds, hands. There were four categories for the (stubby, animate) quadrant: birds, hands, seashells, bowties. To match the number of categories for each quadrant, in the results reported in the main text we select birds and hands in the (stubby, animate) quadrant. In Supplementary Results, we report results when selecting seashells and bowties. To preview, the results are qualitatively similar.

The study was administered online with Pavlovia (https://pavlovia.org/). In each trial, participants were presented with multiple identical target objects of a certain category, along with multiple identical distractor objects of the same category. For each participant, 11 trials were administered for each object category with non-identical target objects. Participants were asked to click on all targets as fast as possible and avoid distractors. The trial ended when all targets had been clicked or when 10 s had passed from the onset of the foraging items, whichever came first.

### Analysis

#### Rank order analysis

##### Within-category consistency

Behavioral preference for each trial was characterized as target clicks second^−1^. For each combination of four categories that covers four quadrants, we performed a split-half analysis across the trials. For each participant and each quadrant, the available trials (#trials = 11) were randomly partitioned into two non-overlapping sets (6 trials and 5 trials). We calculated the rank order of performance across the four quadrants for both sets of trials, ranking by the mean performance of the 6 or 5 trials selected. Then we computed the Spearman’s ρ between the two rank orders. A “shuffled” Spearman’s ρ was computed simultaneously by shuffling the rank order of the 5 trials. This procedure was repeated for 1,000 bootstrap iterations, rendering a distribution of 1,000 ρ and ρ_shuffled_ respectively. Note that for the current study we generally did not employ the Spearman-Brown correction for Spearman’s ρ, as some correlation coefficients can be of negative value.

##### Cross-category consistency

We partitioned the 8 object categories into two non-overlapping sets, each containing one category per quadrant (i.e., four categories). There are 16 combinations of four categories, which translates into 8 possibilities of partitioning the 8 object categories (as partitioning the categories into Set A and Set B is practically the same as partitioning them into Set B and Set A). We correlated the rank order of performance across the two sets of object categories (4 each) using Spearman’s ρ. A “shuffled” Spearman’s ρ was computed simultaneously by shuffling the rank order of one set of 4 object categories. This procedure was bootstrapped across participants for 1,000 iterations, rendering a distribution of 1,000 ρ and ρ_shuffled_ respectively.

#### Landscape analysis

##### Simulation and validation

For the landscape analysis, we segmented the 2D object space (spanning PC1: +/- 30, PC2: +/- 20) into a grid with 2 × 2 “pixels”. For each pixel, we computed the behavioral preference with a partial Spearman’s correlation, based on the center of each pixel:

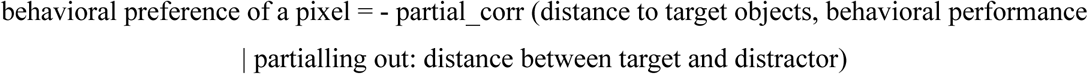

Here we negated the correlation coefficient, as behavioral performance should increase with the decrease of pixel-target distance. The distances were all in Euclidean distance.

To validate the efficacy of our landscape analysis in recovering ground-truth topographies, we performed the following simulations. We defined three ground-truth behavioral performance scenarios: (1) behavioral performance linearly increases along PC1, (2) behavioral performance linearly increases along PC2, (3) behavioral performance linearly increases along the diagonal (by adding the abovementioned two simulations). For each scenario, we simulated a participant performing 1,000 trials, randomly sampling the object space. For each trial, we also generate a “distractor” by randomly sampling the object space. The landscape for each scenario was calculated with the formula above.

##### Within-category consistency

The procedure was identical to the rank order analysis, except that the Spearman’s ρ was calculated across two landscapes.

##### Cross-category consistency

The procedure was identical to the rank order analysis, except that the Spearman’s ρ was calculated across two landscapes.

### Statistical Inference

Statistical significance was determined using non-parametric bootstrap tests. For each analysis, we generated two distributions: a distribution of ρ and a paired shuffled null distribution of ρ_shuffled_. We then derive a new distribution of ρ - ρ_shuffled_. The exact *p*-value for each analysis was calculated as the proportion of values below or above zero (whichever smaller), multiplied by 2 to obtain a two-tailed estimate. Statistical significance was defined with alpha = 0.05.

## SUPPLEMENTARY RESULTS

Here we report the results when we select the categories of bowtie and seashell from the stubby animate quadrant (**Figure S1-2**). Note that bowties and seashells are not animate per se but are animate-looking according to their location in the object space ^3^.

We obtained qualitatively similar results as our selection of eight categories reported in the main text. That is, within-category consistency was generally positive, and cross-category consistency was generally negative.

**Figure S1.**
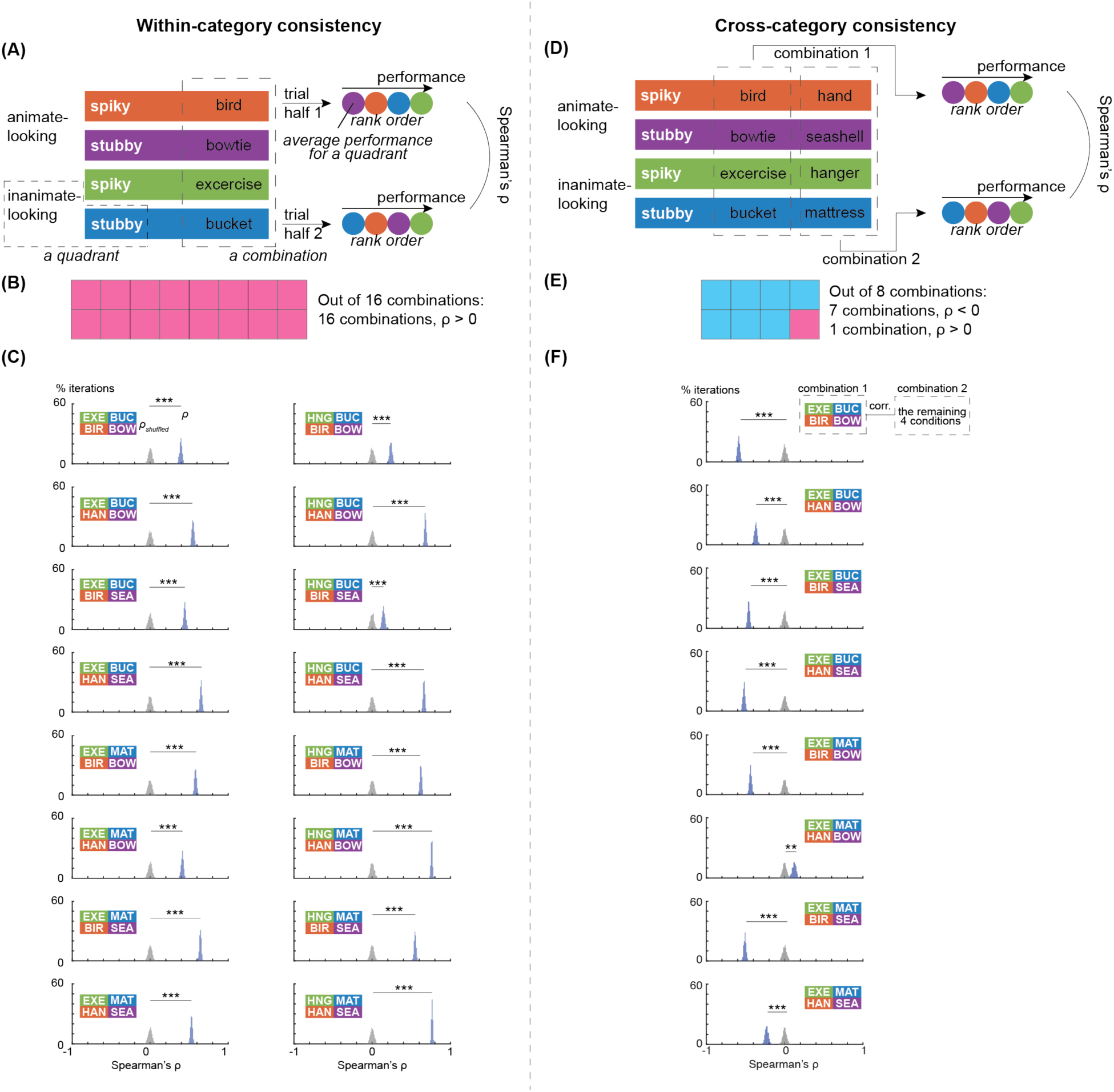
Results of the same analyses from Figure 2 but with a different sampling of eight categories. Abbreviations: EXE: exercise equipment; BUC: bucket; BIR: bird; BOW: bowtie; SEA: seashell; HAN: hand; MAT: mattress; HNG: hanger. ***: *p* < 0.001; **: *p* < 0.01; *: *p* < 0.05.

**Figure S2.**
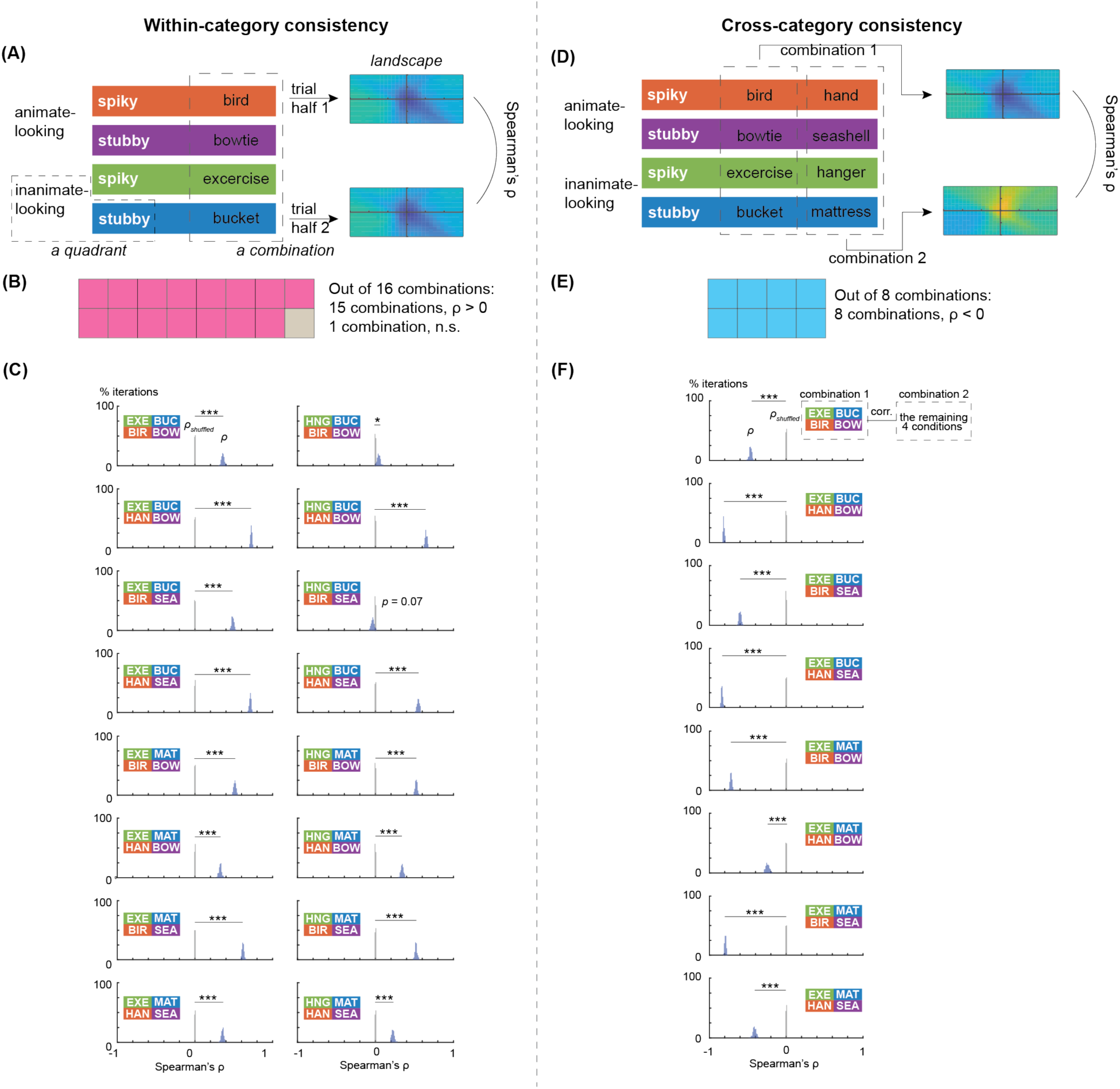
Results of the same analyses from Figure 4 but with a different sampling of eight categories. Abbreviations: EXE: exercise equipment; BUC: bucket; BIR: bird; BOW: bowtie; SEA: seashell; HAN: hand; MAT: mattress; HNG: hanger. ***: *p* < 0.001; **: *p* < 0.01; *: *p* < 0.05.

